# Caffeine ingestion restores morning neuromuscular performance to evening levels in healthy females: A randomised crossover study

**DOI:** 10.1101/2025.10.23.684206

**Authors:** Akshay Singh, Stuart J. Hesketh

**Affiliations:** School of Medicine, University of Lancashire, Preston, UK

## Abstract

**Purpose:** To investigate whether caffeine ingestion offsets the circadian-related decline in morning neuromuscular performance in females.

**Methods:** Thirteen healthy females completed three experimental trials: i) evening placebo (PM_PLAC_), ii) morning placebo (AM_PLAC_), and iii) morning caffeine ingestion (6 mg·kg^−1^; AM_CAFF_). A within-subject, repeated-measures, crossover, single-blind placebo design was used, with all subjects serving as their own controls. Each trial consisted of a maximal voluntary isometric contraction (MVIC) of the knee extensors, followed by intermittent isometric contractions (6 s contraction, 4 s rest) until task failure, and a post-fatigue MVIC. Time to exhaustion, torque and peak force, surface electromyography (RMS amplitude, MDF frequency), rate of perceived exertion, and tympanic temperature were all recorded across conditions.

**Results:** During PM_PLAC_, peak force and time to exhaustion were significantly greater (p < 0.05) than for AM_PLAC_. AM_CAFF_ increased (p < 0.01) peak force in both pre-(+33 %) and post-(+45 %) fatigue MVIC trials compared to PM_PLAC_. Similarly, RMS was increased (p < 0.01) during both AM_CAFF_ and PM_PLAC_ compared to AM_PLAC_, whereas MDF did not change (p > 0.05) across trials. AM_CAFF_ also improved (p < 0.01) time to exhaustion by 43 % compared to AM_PLAC_, and significantly reduced (p < 0.01) rate of perceived exertion, but without any change (p > 0.05) in temperature.

**Conclusions:** Morning ingestion of 6 mg·kg^−1^ of caffeine effectively reverses diurnal reductions in neuromuscular performance of females by restoring peak force and time to exhaustion to evening levels. No change in MDF, but an increased RMS suggests a central rather than peripheral mechanism for caffeine’s ergogenic effects.

## Introduction

Athletic and neuromuscular performance follows a clear diurnal rhythm, with outputs typically lowest in the early morning and peaking in the late afternoon or evening (1-4). This pattern is well documented across strength (1,5), power (6), and endurance tasks (7,8), and reflects the broader influence of circadian regulation on human physiology. Core body temperature, heart rate, blood pressure, and hormonal secretion all fluctuate predictably across the day (9-11), creating time-of-day dependent physiological states that influence readiness for performance. Importantly, these fluctuations have practical implications; for example, athletes competing in morning events may be physiologically disadvantaged compared with later in the day, raising interest in strategies that can offset such declines in performance.

Circadian regulation produces distinct physiological states that influence exercise performance across the day. In the evening, elevated body temperature enhances metabolic reactions and muscle compliance (9,12), while faster nerve conduction and favourable hormonal profiles further facilitate strength, power, and endurance outputs (9,13). Yet, experimental attempts to equalise temperature across time points demonstrate that thermal effects alone cannot fully explain diurnal variations in performance (14-17). Instead, this variation likely arises from the coordinated influence of multiple systems, including neural drive, molecular changes in gene expression, and shifts in substrate metabolism (10,18). Given this complexity, targeted investigation of neuromuscular properties provides a practical lens for understanding how circadian variation influences performance (13,19).

Caffeine is a well-established nutritional strategy with consistent ergogenic effects on exercise performance (20-22). Acting as a non-selective adenosine receptor antagonist, it increases central nervous system excitability, augments neurotransmitter release, and reduces perceived exertion (23,24). Meta-analyses confirm its efficacy across a wide range of tasks, with robust benefits for endurance performance and smaller but meaningful effects on maximal strength and power (25,26). Doses of 3–6 mg·kg^−1^ are most commonly reported to enhance both aerobic and anaerobic outputs, with peak plasma concentrations achieved within 45–60 minutes of ingestion (27,28). Importantly, emerging evidence suggests that caffeine may counteract diurnal decrements in performance, making it a promising strategy to restore morning neuromuscular function to that of evening levels (29,30), thus warranting further investigation.

Despite the general efficacy of caffeine, observed ergogenic effects are heterogeneous and task dependent (31). For example, factors such as exercise modality, training status, dose, timing of ingestion, sex, habitual intake, and methodological factors such as blinding and expectancy (21,32,33) all play a role. Meta-analyses report small-to-moderate average effects, but also highlight considerable between-study variability and a lack of trials directly addressing circadian timing (25,27). These limitations restrict mechanistic inference and further impede the development of population-specific guidance. Sex-specific physiology further complicates interpretation and application. Caffeine metabolism is principally mediated by CYP1A2, whose activity is modulated by sex hormones and menstrual phase, with hormonal contraceptive use known to prolong caffeine clearance and alter plasma exposure, potentially influencing ergogenic efficacy (34,35). Hormonal fluctuations also influence thermoregulation, neuromuscular efficiency, and substrate utilisation, all of which may interact with circadian timing to shape performance. Despite these considerations, most studies remain male-centric, with female participants underrepresented or insufficiently controlled for menstrual status, limiting the applicability of current recommendations (36).

While the ergogenic properties of caffeine are well documented, its capacity to counteract morning declines in neuromuscular performance, particularly in females, remains unclear. The present study therefore aims to determine whether acute caffeine ingestion (6 mg·kg^−1^) can elevate morning neuromuscular performance to evening levels in healthy young females. Specifically, we sought to compare maximal voluntary isometric contraction, time to exhaustion, electromyographic activity, and perceived exertion across morning placebo, morning caffeine, and evening placebo conditions. We hypothesised that (i) neuromuscular performance will be lower in the morning under placebo relative to the evening, (ii) morning caffeine will improve peak force, time to exhaustion, and exertion ratings versus morning placebo, and (iii) increases in performance associated with morning caffeine ingestion will be accompanied by indices of central neural drive, not by alterations in thermoregulation.

## Methods

### Subjects

Thirteen healthy females volunteered to participate in this study (age 24.4 ± 1.1 years; body mass 59.0 ± 10.0 kg; height 159.5 ± 4.4 cm). All participants provided written informed consent prior to enrolment. The study was conducted in accordance with the Declaration of Helsinki, and received approval from the Ethics Committee of the School of Medicine, University of Lancashire (ethics code: 070.24.10.17), and complied with institutional standards for the collection, storage, and confidentiality of human data.

All participants were classified as light caffeine consumers (≤60 mg·day^−1^), based on a validated caffeine consumption questionnaire (37). To minimise confounding influences, participants were instructed to abstain from strenuous exercise outside of testing, as well as from caffeine, alcohol, and smoking for 48 h prior to each trial, given the known effect of nicotine on caffeine metabolism (38). All females were tested in the mid-follicular phase of the menstrual cycle (∼10-15 days), to reduce variability with hormonal fluctuations. Although diet and sleep were not objectively monitored, participants were instructed to refrain from food consumption for at least an hour before each lab visit and to report any irregular sleep patterns over the course of the study. To control for circadian influences, all trials for each participant were conducted at the same time-of-day.

### Study design and supplementation

A within-subject, repeated-measures, crossover, single-blind placebo design was employed, with all participants serving as their own controls. Each participant attended the laboratory to complete three experimental sessions on three consecutive days: (i) evening placebo (17:00; PM_PLAC_); (ii) morning placebo (08:00; AM_PLAC_); and (iii) morning caffeine (08:00; AM_CAFF_; 6 mg·kg^−1^), as shown in Figure 1. This design enabled direct comparison of the main effects of time-of-day (morning vs. afternoon) and caffeine ingestion (0 vs. 6 mg·kg^−1^) on neuromuscular performance. Testing times of 08:00 and 17:00 were selected based on prior evidence demonstrating consistent diurnal variation, with reduced morning (08:00) and enhanced afternoon (17:00) performance (4).

**Figure 1.**
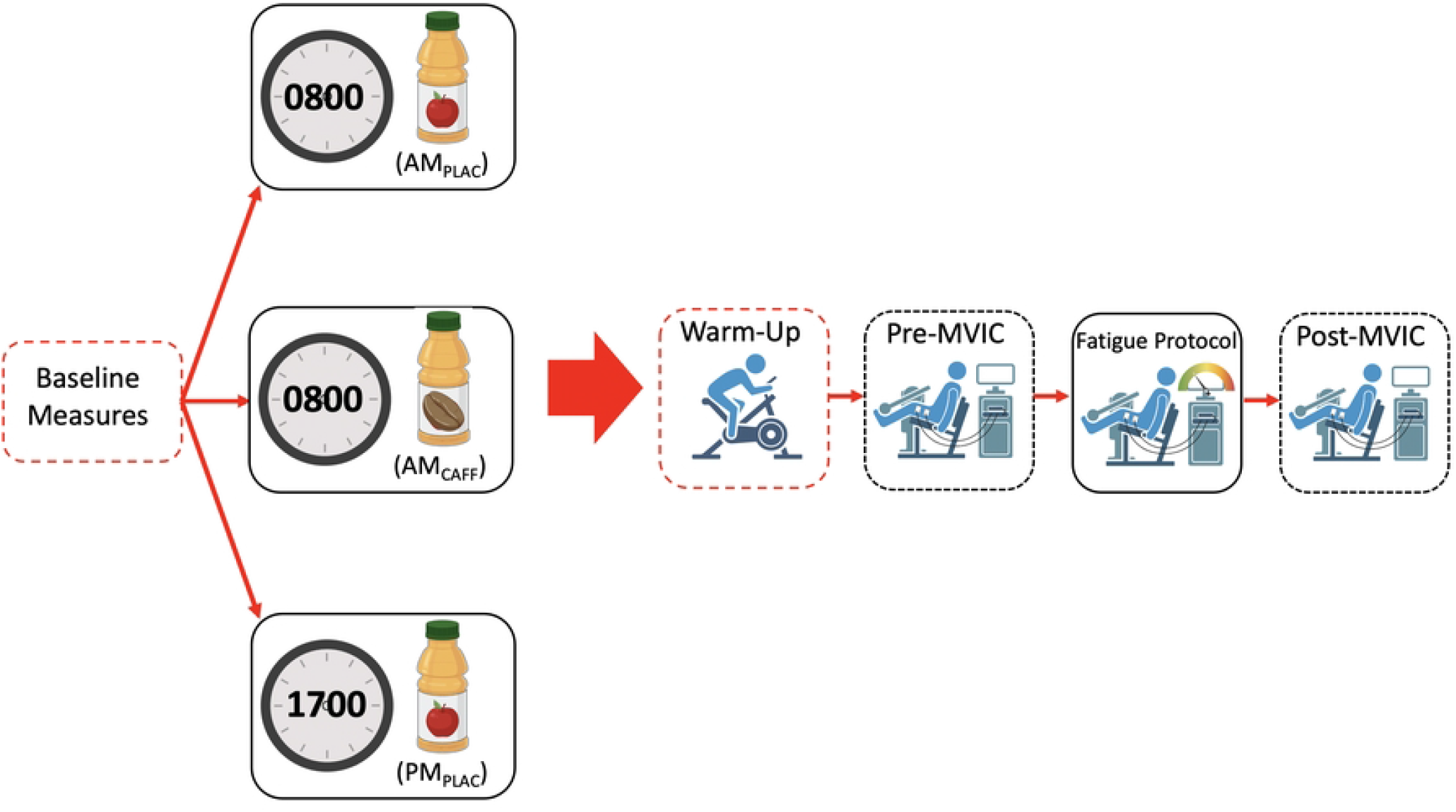
Schematic of experimental design and procedures. Thirteen healthy female participants completed three conditions in a repeated-measures, single-blind, placebo-controlled crossover design. Three trials were completed: morning placebo (AM_PLAC_) at 0800, morning caffeine (AM_CAFF_) at 0800 (6 mg·kg^−1^), and evening placebo (PM_PLAC_) at 1700. Anthropometric measures and core temperature were taken at the beginning of each trial. Each trial consisted of a 5 minute cycling warm-up at 90 rpm, 80 ± 10 W. The first (pre-) maximal voluntary isometric contraction (MVIC) was then performed for 5 seconds at 90 degrees knee flexion with 2 min rest periods between 3 efforts. A fatiguing protocol consisting of 6 second maximal isometric contractions interspersed by 4 seconds rest was then implemented, until peak torque declined to 50 % of pre-MVIC levels, and then post-fatigue MVICs were performed. Surface electromyography and perceived exertion were measured throughout.

A caffeine dose of 6 mg·kg^−1^ body mass was selected, as this range is consistently associated with ergogenic benefits across endurance, strength, and power tasks in controlled trials and meta-analyses (25,27). For the caffeine condition (AM_CAFF_), caffeine powder (ESN caffeine, Germany) was individually weighed for each participant and dissolved in a fruit smoothie consisting of 33 g carbohydrates, 1.6 g protein, 0.15 g fat, and 3.3 g fibre with a total of 143 kcal. In the placebo conditions (AM_PLAC_ and PM_PLAC_), the same fruit drink was provided without caffeine to ensure matched energy content and minimise likelihood of condition detection. Consistent with previous pharmacokinetic studies (39,40), participants ingested the caffeine drink 45 min before testing, expected to coincide with peak plasma concentrations, which typically occur 30-60 min post-ingestion (28,41), thereby maximising the likelihood of testing during peak bioavailability.

### Experimental Protocol

All experimental sessions were conducted in July at the University of Lancashire, Preston, United Kingdom. This period provided a natural circadian window with sunrise at approximately 05:20 and sunset at 22:20. Outdoor temperature averaged 15.5 ± 3.5 °C with relative humidity of 75 ± 5 %. Testing took place in a controlled laboratory environment where ambient temperature was maintained at 25 ± 2 °C to minimise environmental influences on circadian-dependent performance (42).

Following Figure 1, baseline measurements were obtained at the start of each trial, including height (Seca 213 stadiometer, Seca GmbH & Co. KG., Germany), body mass (Seca GmbH & Co. KG., Hamburg, Germany), age, caffeine consumption (37), and tympanic temperature (Braun IRT 4520, Germany). Dominant leg was identified using a validated question-based method described in Van Melick et al, (43). To reduce progressive learning effects, participants completed a familiarisation session during each laboratory visit, replicating the experimental protocol under submaximal conditions. After baseline measurements, to raise muscle temperature and prepare participants for exercise, all individuals performed a standardised warm-up consisting of 5 min of cycling on a stationary ergometer (Keiser M3i, Keiser Corporation, USA) at 90 rpm with a load of 80 ± 10 W.

### Neuromuscular Testing

Surface electromyography (EMG) of the vastus medialis was recorded throughout all testing procedures to measure neuromuscular activation. Prior to electrode placement, the skin was shaved, lightly abraded, and cleaned with an alcohol swab to minimise impedance. Bipolar surface electrodes (Delsys Trigno™ SP-W06, 10 mm interelectrode distance) were positioned according to SENIAM guidelines (44), and the anatomical method described by Rainoldi et al, (45). Electrodes were placed 4 cm above the superior border of the patella, oriented at a 55° angle to the line connecting the anterior superior iliac spine and the patella, aligned with muscle fibres and positioned away from the innervation zone. EMG signals were acquired in real-time using Trigno Discover software, with a band-pass filter of 20–450 Hz applied to minimise noise and motion artefacts before analysis. The EMG output from the vastus medialis was processed using EMGworks Analysis software (v.4.8.0; Delsys Inc., Boston, MA, USA) and full wave rectified. Subsequently, the root mean square (RMS), representing the amplitude of muscle activation, and the median frequency(MDF), obtained via fast Fourier transform as an indicator of motor unit firing behaviour and fatigue, were calculated. A window length of 250 ms with 50 % overlap was applied in accordance with established recommendations (46) to ensure adequate frequency resolution while minimising data dispersion.

Neuromuscular function was assessed using a HUMAC isokinetic dynamometer (HUMAC2015®, Version 15.000.0236; CSMi, Stoughton, MA, USA), calibrated before each session and adjusted to individual anthropometric dimensions. Participants were seated with the dominant leg secured to the lever arm, ensuring alignment of the lateral femoral epicondyle with the axis of rotation. The lower leg was fastened above the malleoli with a padded Velcro strap, and the arms were crossed over the chest to minimise extraneous movement. Positioning was standardised and replicated across sessions.

Maximal voluntary isometric contractions (MVICs) of the knee extensors were performed first (Figure 1). Each trial consisted of a 5-s maximal effort separated by 2 min rest intervals to avoid fatigue (20,47). Three trials were performed, and the highest mean torque value was retained for analysis. Visual feedback and strong verbal encouragement were provided throughout to maximise effort. Following baseline MVICs, participants completed a fatiguing protocol adapted from Pethick and colleagues (47). This consisted of repeated 6-s maximal isometric contractions interspersed with 4-s rest intervals, continued until torque output declined to 50 % of baseline MVIC or volitional failure occurred. The number of completed contractions, time to exhaustion, and ratings of perceived exertion (RPE) were recorded. After a standardised 2-min rest period, participants performed post-fatigue MVICs to assess neuromuscular function under fatigued conditions.

### Statistical analysis

Normality was assessed using Shapiro-Wilk test and all data were found to be suitable for parametric testing. Data are reported as mean ± standard deviation (SD). All statistical analyses were conducted in GraphPad Prism (version 10.5.0; RRID:SCR_002798). One- or two-factor repeated-measures analysis of variance (ANOVA) tests were used to examine within-subject effects of condition and time-of-day. Greenhouse–Geisser corrections were applied, and significant main or interaction effects were followed by Tukey’s post hoc comparisons. Statistical significance was accepted at p < 0.05. Effect sizes were reported using eta-squared (η^2^) and partial eta-squared (ηp^2^).

## Results

Thirteen healthy females completed all experimental trials. Participants were of similar anthropometric stature, 24.4 ± 1.1 years old, with a body mass of 59.0 ± 10.0 kg and height of 159.5 ± 4.4 cm. All were classified as light caffeine consumers (<60 mg·day^-1^) which was recorded using the caffeine consumption questionnaire (37). No adverse events were reported, and all sessions were completed as scheduled. A summary of key outcome measures and statistics across conditions is presented in Table 1 with detailed comparisons below.

**Table 1.**
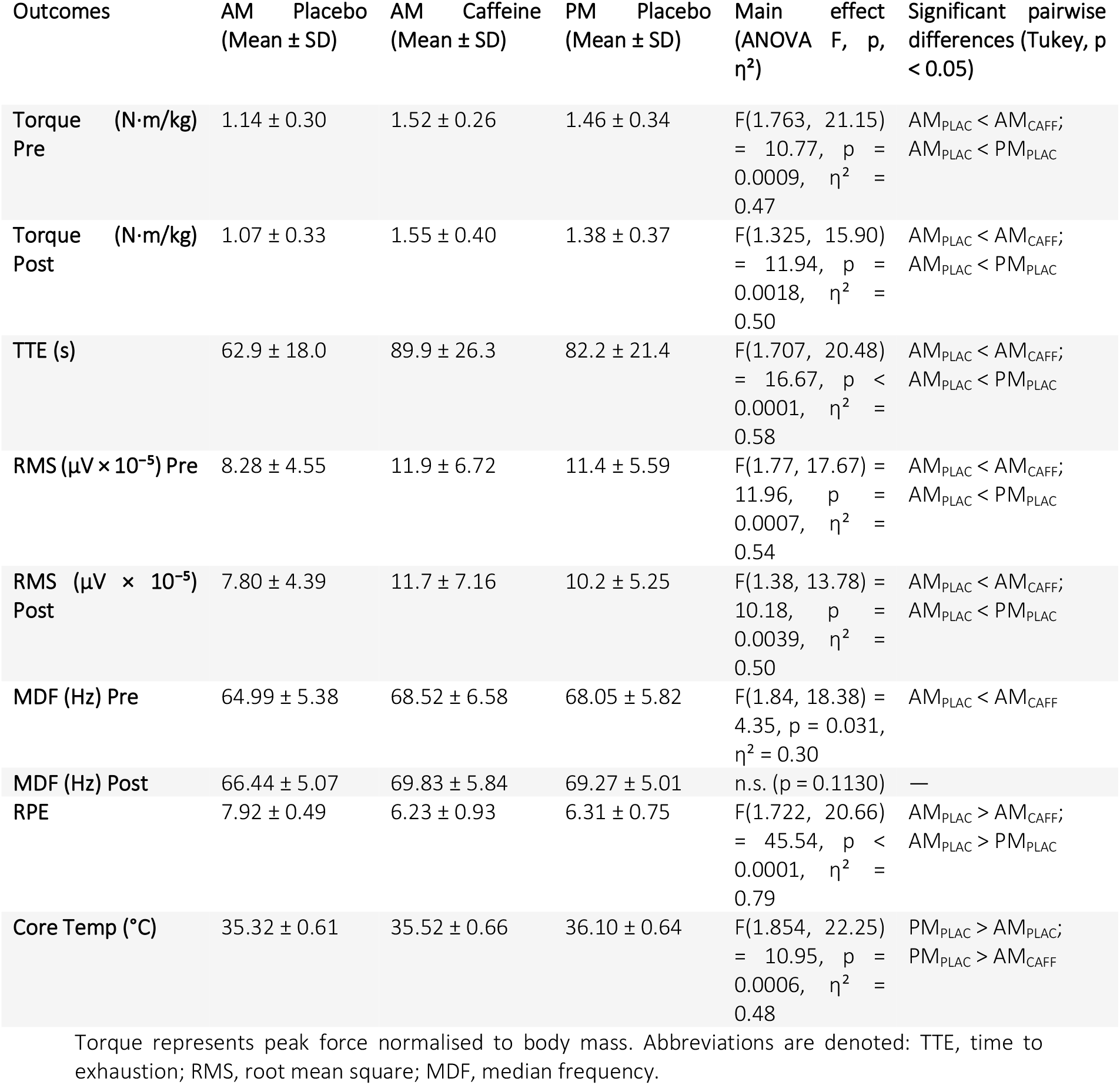
Outcomes measures across experimental conditions.

### Torque

Maximal voluntary torque differed significantly across conditions. Pre-fatigue torque (Figure 2A) showed a main effect of condition, and post-hoc analysis revealed that torque was higher in AM_CAFF_ (1.52 ± 0.26 N·m/kg) and PM_PLAC_ (1.46 ± 0.34 N·m/kg) compared with AM_PLAC_ (1.14 ± 0.30 N·m/kg), representing increases of ∼33 % and ∼28 % respectively (both p < 0.01). A similar pattern was observed post-fatigue (Figure 2B), where post-hoc analysis indicated that torque in AM_CAFF_ (1.55 ± 0.40 N·m/kg) and PM_PLAC_ (1.38 ± 0.37 N·m/kg) was greater than in AM_PLAC_ (1.07 ± 0.33 N·m/kg) by ∼45 % and ∼29 % respectively (both p < 0.01). No differences (p > 0.05) were detected between AM_CAFF_ and PM_PLAC_ at either time point. When pre- and post-fatigue values were analysed together (Figure 2C), torque in AM_PLAC_ remained consistently lower than in both active conditions (all p < 0.01).

**Figure 2.**
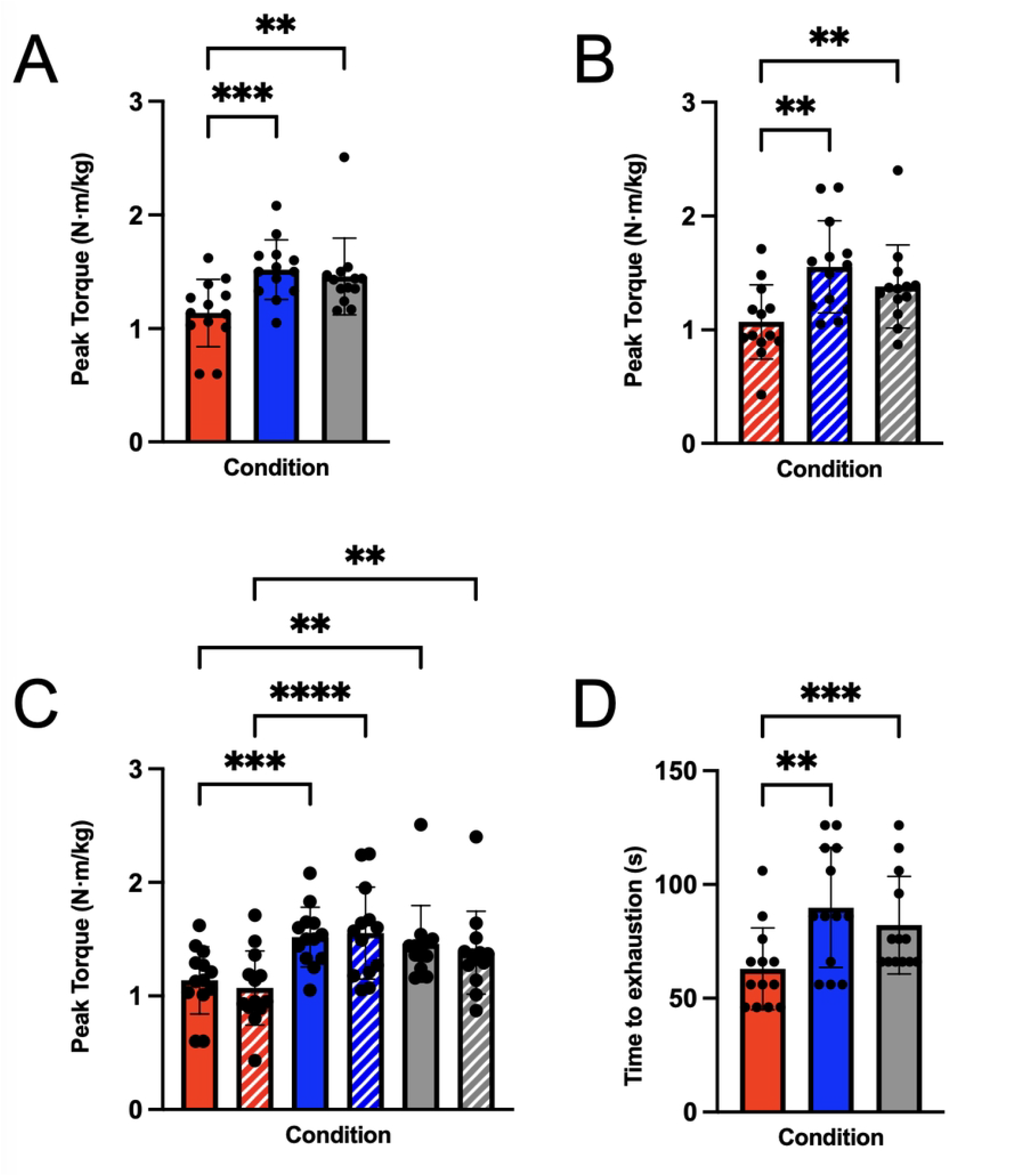
Effect of time-of-day and caffeine ingestion on knee extensor isometric strength. All Data are presented as mean ± SD with individual values plotted. Trials were conducted in the morning (08:00) with placebo (AM_PLAC_), with caffeine ingestion (6 mg.kg^−1^; AM_CAFF_), and in the afternoon (17:00; PM_PLAC_). Red bars represent data for AM_PLAC_ trial, blue bars for AM_CAFF_, and grey bars for PM_PLAC_. All solid bars are for the pre-fatigue MVIC, and hatched bars for the post-fatigue trial. MVIC consisted of 3 times 5 second contractions at 90 degrees flexion with 2-minute rest periods. The fatigue protocol consisted of 6 second maximal contractions interspersed by 4 seconds rest until task failure. (**A**) Peak torque normalised to body mass (N·m/kg) for the pre-fatigue maximal voluntary isometric contraction of the knee extensors. (**B**) Peak torque normalised to body mass (N·m/kg) for the post-fatigue maximal voluntary isometric contraction of the knee extensors. (**C**) Compares pre- and post-fatigue MVICs (**D**) Displays time to exhaustion in seconds for each condition until <50 % of MVIC was attainted or task failure. One-way and Two-way ANOVA was used to assess significance and Tukey’s post hoc testing was applied for multiple comparisons. Significant differences are highlighted for identification, asterisks denote significance level *p < 0.05, **p < 0.01, ***p < 0.001, ****p < 0.0001.

### Time to Exhaustion

Time to exhaustion (Figure 2D) showed a main effect of condition, and post-hoc analysis showed it was longer in AM_CAFF_ (89.9 ± 26.3 s) and PM_PLAC_ (82.2 ± 21.4 s) compared with AM_PLAC_ (62.9 ± 18.0 s), representing increases of ∼43 % and ∼31 % respectively (both p < 0.01). No differences (p > 0.05) were detected between AM_CAFF_ and PM_PLAC_.

### Electromyography

Pre-fatigue root mean square (RMS) (Figure 3A) indicated a main effect and differed significantly across conditions, post-hoc analysis revealed that AM_CAFF_ (11.9 ± 6.7 x 10^-5^ μV) and PM_PLAC_ (11.4 ± 5.6 x 10^-5^ μV) were higher than AM_PLAC_ (8.28 ± 4.6 x 10^-5^ μV), reflecting increases of ∼44 % and ∼38 %, respectively (both p < 0.01). Post-fatigue RMS (Figure 3B) followed the same pattern, with AM_CAFF_ (11.7 ± 7.2 x 10^-5^ μV) and PM_PLAC_ (10.2 ± 5.3 x 10^-5^ μV) exceeding AM_PLAC_ (7.80 ± 4.4 x 10^-5^ μV) by ∼50 % and ∼31 %, respectively (both p < 0.01). No differences (p > 0.05) were measured between AM_CAFF_ and PM_PLAC_ at either time point.

**Figure 3.**
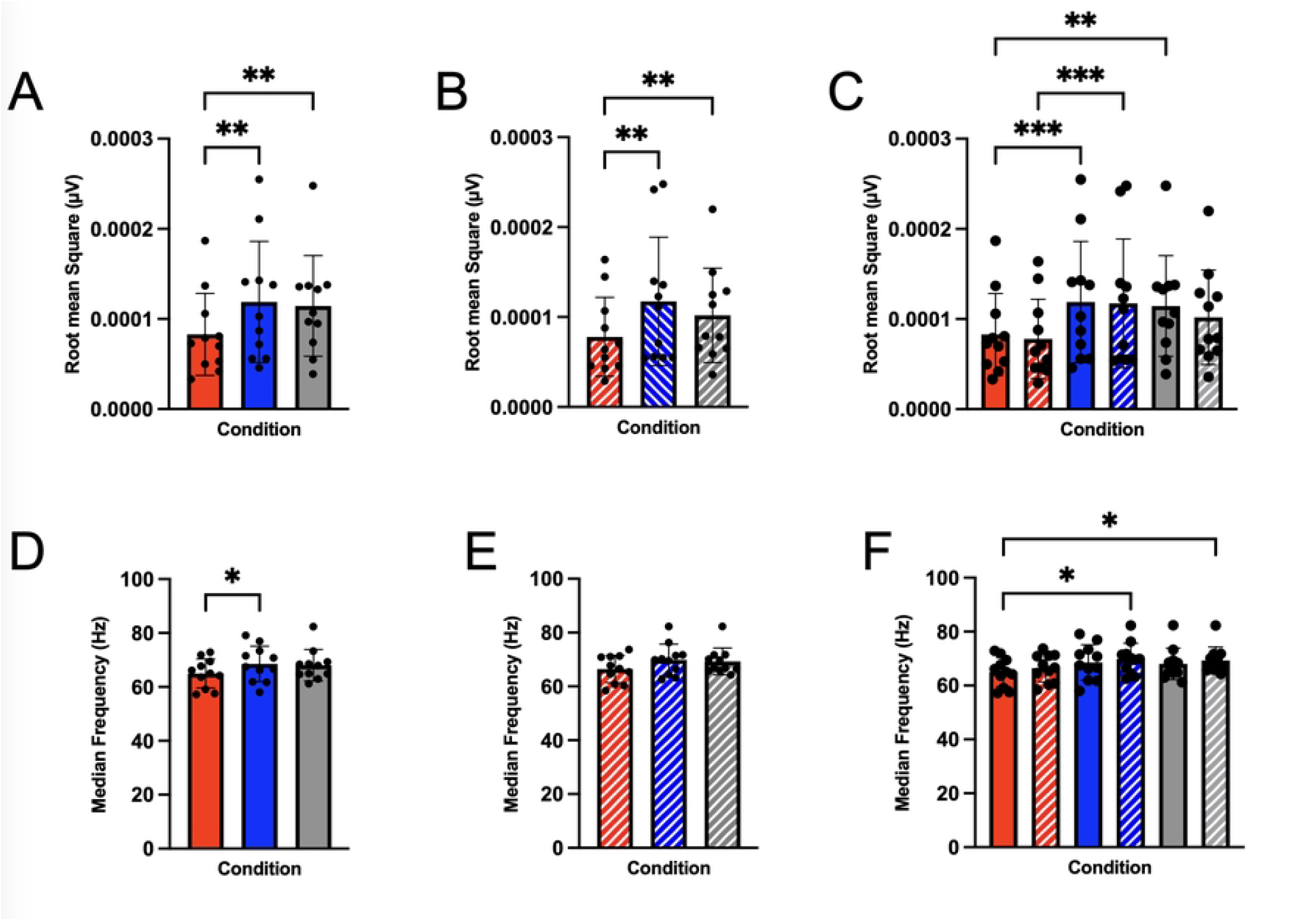
Effect of time-of-day and caffeine ingestion on neuromuscular activation. All data are presented as mean ± SD with individual values plotted. Trials were conducted in the morning (08:00) with placebo (AM_PLAC_), with caffeine ingestion (6 mg.kg^−1^; AM_CAFF_), and in the afternoon (17:00; PM_PLAC_). Red bars represent data for AM_PLAC_ trial, blue bars for AM_CAFF_, and grey bars for PM_PLAC_. All solid bars are for the pre-fatigue MVIC, and hatched bars for the post-fatigue trial. MVIC consisted of 3 times 5 second contractions at 90 degrees flexion with 2-minute rest periods. The fatigue protocol consisted of 6 second maximal contractions interspersed by 4 seconds rest until task failure. Electromyography electrodes were placed on the vastus medialis in the same position for all trials. (**A-C**) Display EMG data for root mean square amplitude for muscle activation during pre- and post-fatigue MVIC measured in µV. (**D-F**) Show EMG data for median frequency for muscle activation during pre- and post-fatigue MVIC measured in Hz. One-way and Two-way ANOVA was used to assess significance and Tukey’s post hoc testing was applied for multiple comparisons. Significant differences are highlighted for identification, asterisks denote significance level *p < 0.05, **p < 0.01, ***p < 0.001, ****p < 0.0001.

Pre-fatigue median frequency (MDF) (Figure 3D) showed a main effect of condition, and post-hoc analysis indicated that AM_CAFF_ (68.5 ± 6.6 Hz) was greater than AM_PLAC_ (65.0 ± 5.4 Hz), representing an increase of ∼5 % (p < 0.05). PM_PLAC_ (68.1 ± 5.8 Hz) did not change (p > 0.05) from either AM_CAFF_ or AM_PLAC_. Post-fatigue MDF (Figure 3E) also did not change across conditions (p >0.05).

### Rate of Perceived Exertion

RPE (Figure 4A) showed a main effect of condition (p < 0.01), and post-hoc analysis revealed that ratings were greater in AM_PLAC_ (7.92 ± 0.49) compared with both AM_CAFF_ (6.23 ± 0.93) and PM_PLAC_ (6.31 ± 0.75), representing a reduction of ∼20 % in both trials (both p < 0.01). No differences (p > 0.05) were measured between AM_CAFF_ and PM_PLAC_.

**Figure 4.**
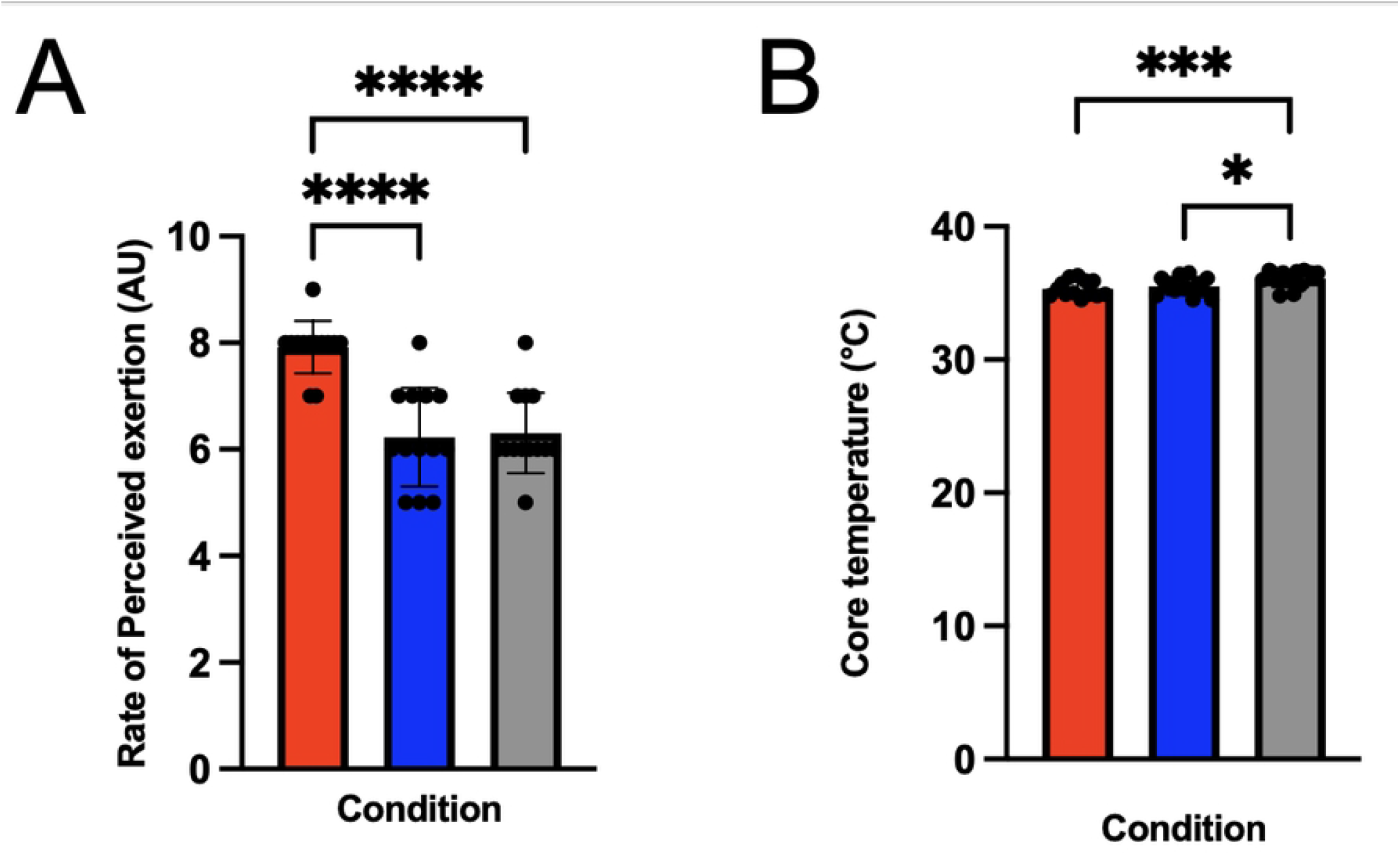
Effect of time-of-day and caffeine ingestion on perception of effort and core temperature. All Data are presented as mean ± SD with individual values plotted. Trials were conducted in the morning (08:00) with placebo (AM_PLAC_), with caffeine ingestion (6 mg.kg^−1^; AM_CAFF_), and in the afternoon (17:00; PM_PLAC_). All trials consisted of 3 MVICs of 5 second contractions at 90 degrees knee flexion with 2-minute rest periods. Red bars represent data for AM_PLAC_ trial, blue bars for AM_CAFF_, and grey bars for PM_PLAC_. (**A**) Rating of perceived effort (scale 1-10) was taken during each trial, and (**B**) tympanic temperature was taken for each measured in degrees Celsius. One-way ANOVA was used to assess significance and Tukey’s post hoc testing was applied for multiple comparisons. Significant differences are highlighted for identification, asterisks denote significance level *p < 0.05, **p < 0.01, ***p < 0.001, ****p < 0.0001.

### Core Temperature

Core temperature (Figure 4B) showed a main effect of condition (p < 0.05), and post-hoc analysis indicated that PM_PLAC_ (36.10 ± 0.64 °C) was almost 3 % higher than both AM_PLAC_ (35.32 ± 0.61 °C) and AM_CAFF_ (35.52 ± 0.66 °C), both p < 0.05. There were no differences (p > 0.05) detected between the two morning conditions.

## Discussion

The present study demonstrates that ingestion 6 mg·kg^−1^ of caffeine 45 minutes before exercise effectively offsets diurnal decline in morning neuromuscular performance in healthy young women. Morning caffeine intake (AM_CAFF_) significantly increased both peak torque by 33–45 %, and time to exhaustion by 43% compared with morning placebo (AM_PLAC_), elevating both outcomes to levels observed in the evening trial (PM_PLAC_) (Figure 2). This improvement was accompanied by higher EMG amplitude (Figure 3), and lower ratings of perceived exertion (Figure 4), while median frequency of EMG and tympanic temperature remained the same (Figure 3 and 4). Our data are consistent with the well-established evidence (1-3,5-7) of diurnal performance differences in physical performance variation in neuromuscular performance, with peak torque and time to exhaustion lower in the morning than in the evening placebo conditions. Collectively, these findings indicate that caffeine reverses morning decrements in neuromuscular performance, primarily via central mechanisms rather than peripheral or thermoregulatory processes.

Previous investigations of diurnal differences in performance have attempted to attribute daily fluctuations in performance to core temperature (2,14-17), metabolic activity (48), and neural function (49,50). We investigated the effect of ergogenic aid consumption and associated effects of core temperature and neuromuscular function. Importantly, our data show that in females caffeine ingestion can elevate morning performance to evening levels, in agreement with previous work demonstrating that caffeine mitigates circadian-related decrements in strength and power of male athletes (29,30). Meta-analyses further highlight consistent ergogenic effects across both strength and endurance tasks (25,26,51), supporting the robustness of the current findings. For example, the present study adds new evidence that this effect also occurs in females, suggesting the mechanism is centrally mediated and independent of temperature.

Not all studies have observed uniform effects. Some evidence suggests that caffeine exerts greater benefits in tasks involving larger muscle groups (31), or dynamic contractions (52). For example, Mora-Rodríguez et al, (29) reported improvements in dynamic but not isometric contractions at 3 mg·kg^−1^, whereas benefits at 6 mg·kg^−1^ were evident in maximal squats, but not bench press (30). In contrast, our study demonstrates clear improvement in isometric torque of the knee extensors, likely reflecting both the involvement of a large lower-limb muscle group and the specific demands of sustained maximal activation. This adds to the literature by showing that caffeine can also enhance isometric performance under fatigue when sufficient dose and muscle mass engaged.

The absence of a direct association between core temperature and performance provides mechanistic insight. As expected, afternoon trials were accompanied by elevated temperature, consistent with circadian regulation (9). Yet, caffeine improved torque and time to exhaustion in the morning without increasing temperature, suggesting that thermoregulatory changes were not responsible for the ergogenic effect in our data. This further aligns with previous work showing caffeine can improve neuromuscular performance independent of body temperature (29,30), and supports a centrally mediated mechanism now demonstrated for the first time in females.

Instead, our EMG findings suggest enhanced central neural drive. RMS amplitude, an index of motor unit recruitment and activation, was higher with caffeine ingestion, while MDF, a spectral index related to motor unit firing behaviour and fatigue, remained unchanged. This pattern indicates more sustained motor unit activation rather than altered peripheral conduction, in line with earlier reports showing caffeine-induced increases in EMG amplitude without significant changes in spectral fatigue markers (20,24,46). These results are consistent with the established central actions of caffeine as an adenosine receptor antagonist, which increases cortical excitability, augments neurotransmitter release, and reduces perceived exertion (53,54). The significant reduction in RPE observed here further supports this central mechanism, consistent with reports that caffeine lowers perception of effort and enhances tolerance to fatigue (7,8,46,55).

A critical contribution of this study is its focus on females, a population historically under-represented in caffeine research (36). Sex hormones influence caffeine metabolism via modulation of CYP1A2 activity, with menstrual phase and contraceptive use known to affect clearance rates and plasma exposure (34,35). Hormonal fluctuations also influence thermoregulation, substrate metabolism, and neuromuscular efficiency. Despite these complexities, most caffeine research has been male-centric or insufficiently controlled for female-specific physiology. By recruiting healthy young females and partially controlling for menstrual cycle timing, the present study addresses this gap and provides evidence that caffeine rescues morning neuromuscular performance in this group. Future studies should expand on this work to older or post-menopausal women, where reduced oestrogen and progesterone levels may alter the pharmacokinetics and efficacy of caffeine (56,57).

Several limitations warrant consideration. First, while participants refrained from caffeine for 48 hours prior to testing, habitual caffeine intake was not fully quantified, leaving potential residual effects. Chronotype and sleep were not rigorously controlled, each of which can modulate circadian physiology and caffeine metabolism (58,19). Third, the single-blind design carries risk of expectancy bias, as the physiological side effects of caffeine may have reduced masking integrity (33). Plasma caffeine concentrations were also not measured, preventing direct verification of absorption kinetics. Finally, EMG provides indirect insight into neuromuscular activation, but more definitive approaches such as interpolated twitch techniques or transcranial magnetic stimulation would better disentangle central versus peripheral contributions (59).

Despite these limitations, the findings offer important contributions to both theory and practice. Theoretically, these data support the notion that the ergogenic potential of caffeine in the morning operates primarily through central rather than thermal or peripheral pathways. Practically, they suggest two strategies to mitigate diurnal performance decrements i) scheduling training or competition in the afternoon to align with circadian peaks, or ii) ingesting caffeine (6 mg·kg^−1^) 45-60 minutes before morning exercise. Given inter-individual variability in caffeine sensitivity, practitioners should tailor dosing (3-6 mg·kg^−1^) based on tolerance and habitual intake, while monitoring for side effects.

## Conclusion

In summary, acute caffeine ingestion (6 mg·kg^−1^) taken 45 minutes before exercise improves morning neuromuscular performance in young females to that of evening levels. Improvements in torque, time to exhaustion, EMG amplitude, and perceived exertion occurred without changes in EMG frequency characteristics or core temperature, indicating a central rather than peripheral or thermoregulatory mechanism. These findings extend existing literature by demonstrating for the first time that caffeine effectively offsets circadian-related performance decrements in females, a group under-represented in scientific research.

## Availability of Data and Materials

The data that support the findings of this study are available upon reasonable request from the corresponding author.

